# IL-7R signalling activates widespread V_H_ and D_H_ gene usage to drive antibody diversity in bone marrow B cells

**DOI:** 10.1101/2020.09.24.298000

**Authors:** Amanda Baizan-Edge, Bryony A. Stubbs, Michael J. T. Stubbington, Daniel J. Bolland, Kristina Tabbada, Simon Andrews, Anne E. Corcoran

**Affiliations:** Nuclear Dynamics Programme, Babraham Institute, Babraham Research Campus, Cambridge CB22 3AT, UK; Lymphocyte Signalling and Development Programme, Babraham Institute, Babraham Research Campus, Cambridge CB22 3AT, UK; Bioinformatics Group, Babraham Institute, Babraham Research Campus, Cambridge CB22 3AT, UK; Centre for Gene Regulation and Expression, School of Life Sciences, University of Dundee, Dow Street, Dundee DD1 5EH, UK; M.J.T.S. is an employee, share- and option-holder of 10x Genomics, 10x Genomics, 6230 Stoneridge Mall Road, Pleasanton, CA 94588, USA

**Keywords:** B cell development, Igh recombination, IL-7R signalling, bone marrow, fetal liver, pro-B cells, non-coding transcription, accessibility

## Abstract

Generation of the primary antibody repertoire requires V(D)J recombination of hundreds of gene segments in the immunoglobulin heavy chain (Igh) locus. It has been proposed that interleukin-7 receptor (IL-7R) signalling is necessary for Igh recombination, but this has been challenging to partition from the receptor’s role in B cell survival and proliferation. By generating the first detailed description of the Igh repertoire of murine IL-7Rα^-/-^ bone marrow B cells, we demonstrate that IL-7R signalling profoundly influences V_H_ gene selection during V_H_-to-DJ_H_ recombination. We find skewing towards usage of 3’ V_H_ genes during *de novo* V_H_-to-DJ_H_ recombination that is more severe than the fetal liver (FL) B cell repertoire, and we now show a role for IL-7R signalling in D_H_-to-J_H_ recombination. Transcriptome and accessibility analyses suggests reduced expression of B lineage-specific transcription factors (TFs) and their targets, and loss of D_H_ and V_H_ antisense transcription in IL-7Rα^-/-^ B cells. These results refute models suggesting that IL-7R signalling is only required for survival and proliferation, and demonstrate a pivotal role in shaping the Igh repertoire by activating underpinning epigenetic mechanisms.

## Introduction

Interleukin-7 (IL-7) is a critical cytokine for B and T lymphocyte development. It is bound by the IL-7 receptor (IL-7R), composed of the common gamma chain (γc), shared with the IL-2R, IL-15R and others, and the IL-7-specific IL-7Rα chain (CD127 - encoded by *Il7r*), which can also pair with the thymic stromal lymphopoietin receptor (TSLPR), important in fetal liver (FL) pro-B and pre-B cell survival (Vosshenrich et al., 2003; Rochman et al., 2009). Binding of IL-7 to the IL-7R in pro-B cells activates several signalling pathways, including Janus kinase/signal transducers and activators of transcription (JAK/STAT), phosphoinositide-3 kinase (PI3K)/AKT, MAPK/ERK, and PLCγ/PKC/mTOR, and has been associated with proliferation and survival, B cell lineage commitment and Igh recombination (Corcoran et al 1996; Corcoran 1998; reviewed by Corfe et al., 2012; Yu et al., 2017).

IL-7R signalling is essential for B cell commitment. Its absence in IL-7Rα^-/-^ mice impairs early B cell development from the common lymphoid progenitor (CLP) stage, resulting in reduced B lineage progenitors and impaired B-lineage potential (Peschon et al., 1994; Miller et al., 2002; Erlandsson et al., 2004; Dias et al., 2005, Medina et al., 2013). This is due in part to failure to activate EBF1, a crucial TF for B lineage specification (Dias et al., 2005; Kikuchi et al., 2005; Tsapogas et al., 2011; Thal et al., 2009; Pongubala et al., 2008; Roessler et al., 2007; Boller and Grosschedl, 2014). PAX5, a key TF for B cell commitment (Nutt et al; 1999; Fuxa et al., 2007; Rumfelt et al., 2006), is also reduced in IL-7Rα^-/-^ pro-B cells (Corcoran et al., 1998), but since EBF1 contributes to the activation of PAX5 (O’Riordan and Grosschedl 1999; Decker et al., 2009; Lin et al., 2010), this may be a downstream consequence of reduced EBF1 expression. However, the block in B cell development in IL-7Rα^-/-^ mice is not complete, with some cells progressing to the pre-pro-B stage (Kikuchi et al., 2005; Peschon et al., 1994; Miller et al., 2002), and fewer cells progressing to the CD19^+^ pro-B compartment (Corcoran et al., 1998; Miller et al., 2002).

Although IL-7Rα^-/-^ B cells in the bone marrow (BM) undergo Igh V_H_-DJ_H_ recombination (Corcoran et al., 1998), a pro-B-specific feature, their usage of V_H_ genes is severely restricted, indicating a role of IL-7R signalling in this process. Importantly, this role is independent of proliferation: IL-7Rα^-/-^ cells expressing an IL-7Rα chain lacking Tyr449, required for PI3K signalling, expressed a rearranged heavy chain polypeptide (Igμ), indicating they had undergone V(D)J recombination, but failed to proliferate *in vitro* (Corcoran et al., 1996). Conversely, a chimeric receptor comprising the extracellular domain of the IL-7Rα chain fused to the intracellular domain of the IL-2Rβ chain restored proliferation in IL-7Rα^-/-^ cells, but not Igμ expression, indicating a non-redundant role for the IL-7R in Igh V(D)J recombination.

A diverse antibody repertoire requires inclusion of all the available V_H_ and D_H_ genes. Many features influence the choice of V_H_ genes for V(D)J recombination. Large-scale processes, involving sense and antisense non-coding transcription, histone modifications and Ig locus contraction by DNA looping, are thought to increase the accessibility of distal 5’ V_H_ genes to the recombination centre over the DJ region, where the Recombination Activating Genes (RAG) 1 and 2 associate, thereby facilitating equal spatial proximity for V to DJ recombination (Ji et al., 2010, Corcoran et al., 1998; Bolland et al., 2004; Yancopoulos and Alt, 1985, Chowdhury and Sen, 2003; Fuxa et al., 2004; Jhunjhunwala et al., 2008; Sayegh, 2005; Stubbington and Corcoran, 2013). Nevertheless, V_H_ genes recombine at widely different frequencies (Bolland et al., 2016). Additionally, frequently recombining V_H_ genes have one of two local, mutually exclusive, active chromatin states close to the recombination signal sequence (RSS) 3’ of V_H_ genes: the enhancer state, characterised by the active histone modification H3K4me1 and PAX5, IRF4, and YY1 binding, and the architectural state, defined by CTCF and RAD21 binding (Bolland et al., 2016).

IL-7Rα^-/-^ pro-B cells *in vivo* displayed decreased germline (non-coding) transcription and recombination of 5’ V_H_ genes (Corcoran et al., 1998), prompting the hypothesis that the IL-7R may influence Igh recombination through increasing the accessibility of 5’ V_H_ genes. This hypothesis was supported by subsequent studies linking IL-7R signalling and active histone modifications. First, histone H3 hyperacetylation occurs over the 5’ V_H_-J558 genes in an IL-7-dependent manner (Chowdhury et al., 2001; Johnson et al., 2003). Second, IL-7^-/-^ pro-B cells exhibited reduced H3K36me2 across the Igh locus, whilst transgenic overexpression of IL-7 increased H3K36me2 deposition (Xu et al., 2008). Third, STAT5 knockdown experiments showed that STAT5 regulates germline transcription, histone acetylation and DNA recombination of the 5’ V_H_ gene segments, suggesting that IL-7R activation of the Igh locus is mediated by its activation of STAT5 (Bertolino et al., 2005). The IL-7R has also been shown to inhibit Rag recombinase expression. Apparent conflict between this and its role in proliferation, versus a role in Igh accessibility were resolved by demonstration that heterogeneous expression of the IL-7R during the cell cycle regulates Rag gene expression to prevent DNA breaks during replication, while Igh locus accessibility is maintained throughout the cell cycle (Johnson et al 2012).

However, the link between Igh accessibility and IL-7R signalling has been challenged. Conditional deletion of STAT5 from the CLP stage resulted in a reduction of all later B cell stages in a manner similar to IL-7Rα^-/-^ mice (Malin et al., 2010). Cells rescued by the pro-survival factor BCL-2 exhibited no deficiency in PAX5 expression or 5’ V_H_ recombination, and the authors concluded that the dominant role of STAT5 was pro-B cell survival. However, IL-7Rα^-/-^ B cells cannot be rescued by Eu-BCL-2 expression (Maraskovsky et al., 1998), and only partially by vav-BCL2 (Malin et al., 2010), indicating that the IL-7R has additional downstream signalling mechanisms and roles in B cell development beyond STAT5 activation and survival. Furthermore, the IL-7R Y449F mutant that abrogates both STAT5 and PI3K activation nevertheless enables recombination, indicating that an alternative IL-7R-related pathway is at play (Corcoran et al., 1996).

Other reports have argued that the lack of IL-7Rα prevents B cell progression beyond the pre-pro-B stage, and that any committed B cells in the BM originate from the FL (Kikuchi et al., 2005; Peschon et al., 1994; Miller et al., 2002; Carvalho et al., 2001). These models were supported by similarity in V_H_ repertoire between IL-7Rα^-/-^ and FL B cells, both preferentially recombining 3’ V_H_ genes (Corcoran et al., 1998; Jeong and Teale, 1988; Yancopoulos et al., 1988). However, definitive conclusions have been hampered by incomplete knowledge of the V_H_ genes of the Igh locus, provided at a later date (Johnston et al., 2006), and low-resolution Igh repertoire assays.

With next generation sequencing (NGS) technologies enabling more in-depth and comprehensive analysis of Igh repertoires and gene expression, we have re-visited the IL-7Rα^-/-^ mouse strain (Peschon et al., 1994) to resolve the uncertainties described above, which confound a clear and comprehensive picture of the role of the IL-7R in B cell development. Using VDJ-seq (Bolland et al., 2016), an unbiased DNA-based NGS technique that captures Igh DJ_H_ and VDJ_H_ recombined sequences, we have fully characterised the Igh repertoire in BM B cells of IL-7Rα^-/-^ mice. We have found that Igh recombination takes place, but is defective from the CLP stage onwards in IL-7Rα^-/-^ mice, since widespread use of gene segments in both D_H_-J_H_ (which begins in CLPs and finishes in pre-pro-B cells) and V_H_-D_H_ recombination (occurring in pro-B cells) was severely impaired by the lack of IL-7R signalling.

Importantly, we have also revealed that IL-7Rα^-/-^ BM B cells expressing a reduced antibody repertoire are not simply derived from the FL. First, a defining feature of BM V(D)J recombination is the expression of terminal deoxynucleotide transferase (TdT), responsible for junctional insertion of N-nucleotides (Feeney et al., 1990; Li et al., 1993). Junctions between V_H_, D_H_ and J_H_ gene segments in IL-7Rα^-/-^ BM B cell recombined Igh loci, similar to WT, showed substantially increased variability compared with FL B cell recombined sequences, indicating that BM pro-B cells in IL-7Rα^-/-^ mice undergo de *novo* V(D)J recombination. Second, IL-7R^-/-^ BM B cells fail to use a large proportion of 5’ V_H_ genes, exhibiting a much more severe reduction in repertoire diversity than FL. Transcriptome analysis revealed loss of large-scale antisense intergenic transcripts in both the D_H_ and V_H_ regions and reduced expression of key transcription factors required for Igh recombination, including EBF1 and Pax5, while local V_H_ gene accessibility was unaltered. These findings, together with previous studies, demonstrate that IL-7R signalling specifically promotes Igh repertoire diversity in BM pro-B cells by activating several underpinning epigenetic mechanisms that enable large-scale access to V_H_ and D_H_ genes.

## Results

### Usage of V and D genes in Igh recombination is severely skewed in pro-B cells lacking the IL-7Rα chain

VDJ-seq was performed on two IL-7Rα^-/-^ BM and two WT FL pro-B cell samples and compared with two WT BM replicate datasets previously generated in our lab (GEO: GSE80155), (Bolland et al. 2016). Replicate libraries for all genotypes were highly correlated indicating VDJ-seq is highly reproducible (Figure S1). The ratio of VDJ_H_ to DJ_H_ recombined products in IL-7Rα^-/-^ was similar to WT, indicating there was no gross defect in V(D)J recombination, i.e. dynamic progression through first D_H_ to J_H_, followed by V_H_ to DJ_H_ recombination was not slowed (Figure S2). A binomial test was applied to determine genes with significantly greater read counts than expected by chance, which were considered to be actively recombining (Bolland et al., 2016) (S Table 1). Consistent with a requirement for IL-7R signalling to enable participation of V_H_ genes in recombination, fewer V_H_ genes passed the binomial test in IL-7Rα^-/-^ (84 genes) relative to WT pro-B cells (128 genes). All but three of these 84 V_H_ genes were within the group of 128 recombining V_H_ genes in WT (S Table 1).

To visualise V_H_ gene recombination frequencies and compare between genotypes, V_H_ gene quantifications were normalised to the total number of reads over all V_H_ genes for each genotype (Figure 1A). Analysis of V_H_ gene frequency confirmed and expanded previous low-resolution reports on two V gene families in the IL-7Rα^-/-^ V_H_ repertoire (Corcoran et al., 1998), showing a much higher proportion of sequences mapped to the most 3’ V_H_ genes than in the WT repertoire (Figure 1B). In particular, the first five V_H_ genes comprised 45% of the total repertoire, compared with 20% in WT. Of the 44 V_H_ genes that recombine in WT, but were not actively recombining in IL-7Rα^-/-^ BM, the vast majority were at the 5’ end of the V region, including several that normally recombine at high frequency, including J558.16.106, J558.26.116, J558.67.166.

**Figure 1:**
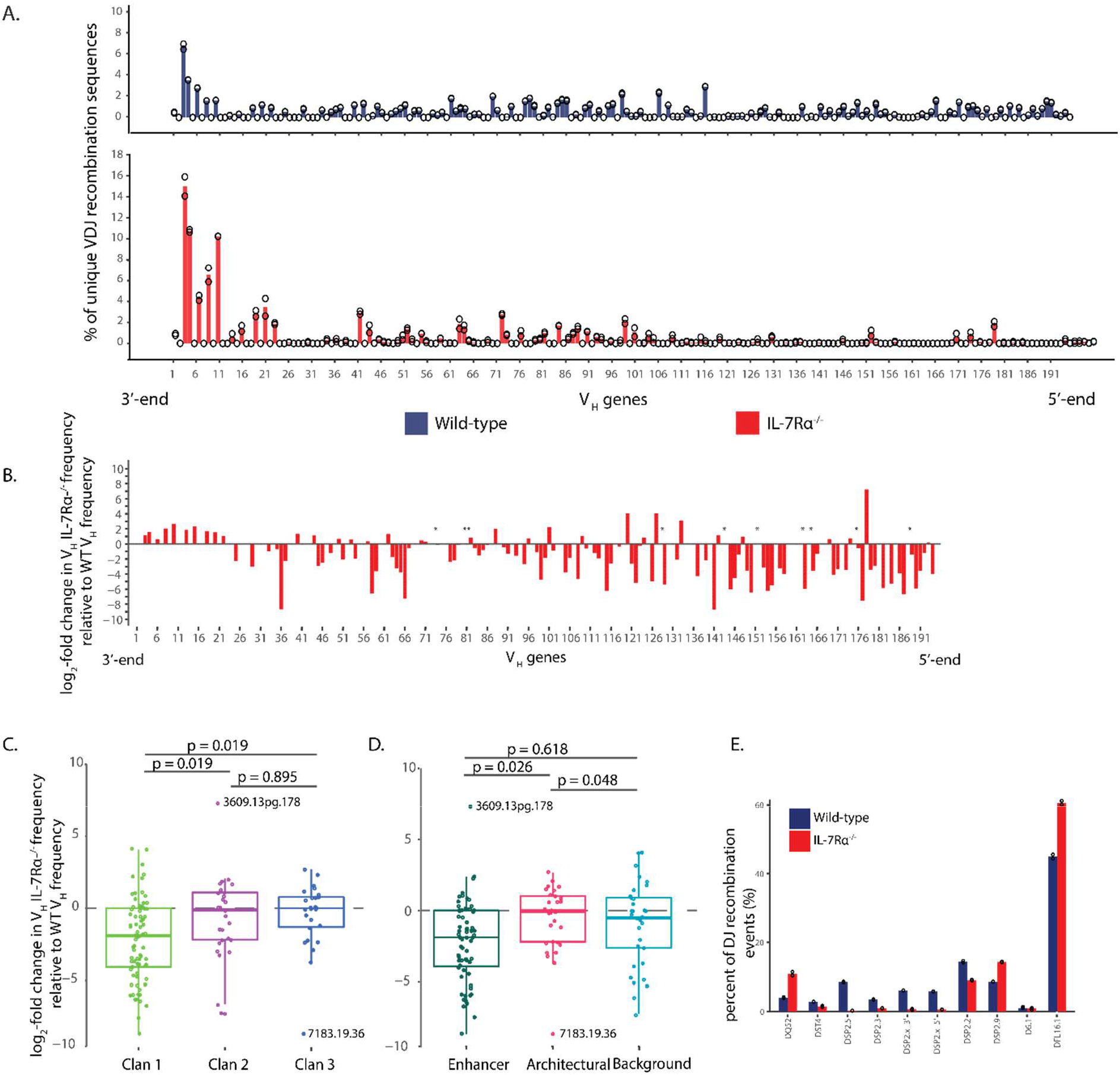
IL-7Rα^-/-^ pro-B cells have impaired V_H_ and D_H_ repertoires. A. Recombination frequencies of the 195 V_H_ genes measured by VDJ-seq, for BM pro-B cells from WT (blue), and IL-7Rα-/- (red) mice (20 IL-7Rα-/- per replicate). Two replicates were generated and are shown as open circles (height of bar represents average). Reverse-strand reads were quantified for each V_H_ gene and shown as a percentage of total number of reads quantified. For V_H_ gene number legend see supplementary table 1. B. The mean of each V_H_ gene reads in two libraries were divided with the WT replicate mean value followed by log2-transformations. Only genes that were recombining in either genotype are shown. * represents V_H_ genes that had value 0 in IL-7Rα-/- (only occurring in WT replicates). For list of V_H_ genes with raw read counts and recombining/non-recombining classification, see supplementary table 1. Log2 values for each gene in graph B (excluding those marked by *) were grouped by C. evolutionary origin: clan 1 (n = 78), clan 2 (n = 27) and clan 3 (n = 26), ANOVA (degrees freedom = 2, F-value = 5.39, p-value = 0.005); D. chromatin state: enhancer (n = 68), architectural (n = 30) and background (n = 33); ANOVA (degrees freedom = 2, F-value = 4.54, p-value = 0.012) E. Reads were quantified for each D_H_ gene and shown as a percentage of total number of reads quantified for two replicates of WT (blue), and IL-7Rα-/- (red) pro-B cells.

The 195 V_H_ genes segregate into 16 families within 3 clans that have evolved separately (Johnston et al., 2006). The evolutionary origin of V_H_ genes correlates with differential TF binding and chromatin state around V_H_ genes (Bolland et al., 2016). Since accessibility, TF expression and chromatin state have been linked to IL-7R signalling, we investigated the relationship between recombination frequency and V_H_ gene clan, or chromatin state in IL-7R^-/-^ pro-B cells. Overall, V_H_ genes in clan 1 recombine at lower frequency in the IL-7Rα^-/-^ samples relative to WT, while clan 2 and 3 V_H_ genes recombine at similar frequencies to WT (Figure 1C). When the data are broken down to V_H_ gene families, a more nuanced picture emerges. Within clan 2 and 3, 7183 and Q52, the two largest and most 3’ V_H_ families, recombine more frequently in IL-7Rα^-/-^. However, several of the smaller families in the middle region recombine less frequently (Figure S3). The enhancer state V_H_ genes (including those from clan 1 and the distal 3609 family from clan 2) were significantly less frequently recombined overall relative to the architectural state V_H_ genes (clan 2, except 3609, and clan 3 - Figure 1D). Nevertheless, some of the architectural state V_H_ families also recombined less frequently. Together these data suggest that loss of IL-7R impacts on V_H_ genes in the enhancer state (distal and middle genes) and on several middle region families in the architectural state. This distribution also applies to the clans: loss of IL7R reduces recombination of Clan 1 (mostly distal V genes) as well as the middle genes from clans 2 and 3. Importantly this suggests that the IL-7R does not influence either clans or chromatin states selectively, but rather linear positioning in the Igh V region i.e. loss of IL-7R impairs recombination of middle and 5’ V genes in the Igh locus.

Actively recombining D_H_ genes were also identified by binomial test. Although previous reports did not show a reduction in overall D-J recombination (Corcoran et al., 1998; Bertolino et al., 2005), our detailed analysis with VDJ-seq revealed profound changes in individual D_H_ usage in IL-7Rα^-/-^ pro-B cells. Several DSP D_H_ gene segments (DSP2}5’, DSP2}3’, DSP2.3 and DSP2.5) that were actively recombined in WT were not represented in the IL-7Rα^-/-^ repertoire (Figure 1E), indicating that these centrally positioned D_H_ gene segments do not recombine in IL-7Rα^-/-^ pro-B cells. Conversely, relative recombination frequencies of the most 3’ D_H_ gene, DQ52, the most 5’, DFL16.1 and its adjacent DSP, were increased in IL-7Rα^-/-^ cells.

### IL-7Rα^-/-^ pro-B cells do not originate from the FL

A proximity bias in usage of V_H_ genes similar to that observed in IL-7Rα^-/-^ cells was previously reported in low resolution studies in FL pro-B cells (Jeong and Teale, 1988; Yancopoulos et al., 1988). Therefore, we investigated whether recombination events in IL-7Rα^-/-^ BM were comparable to those generated by B cells derived from FL, rather than *de novo* in the BM, as previously suggested (Carvalho et al., 2001). VDJ-seq was performed on WT embryonic day 15.5 pro-B cells, and the data compared with sequences derived from the BM of IL-7Rα^-/-^ and WT mice. Notably, the ratio of DJ to VDJ recombinants in FL was 12:1, in contrast to WT and IL-7Rα^-/-^ BM ratios of approximately 2:1 and 1:1, respectively (Figure S2).

Consistent with previous reports, the FL V_H_ gene repertoire exhibited a 3’ bias relative to WT, including more frequent use of the most recombined V_H_ gene, V_H_-81X (11% compared with 7% for WT BM) (Figure 2A and S4). However, IL-7Rα^-/-^ cells showed a more pronounced phenotype, with V_H_-81X comprising 14% of VDJ recombined sequences. Additionally, many 5’ V_H_ genes recombined less frequently than in FL pro-B cells (Figure 2A).

**Figure 2:**
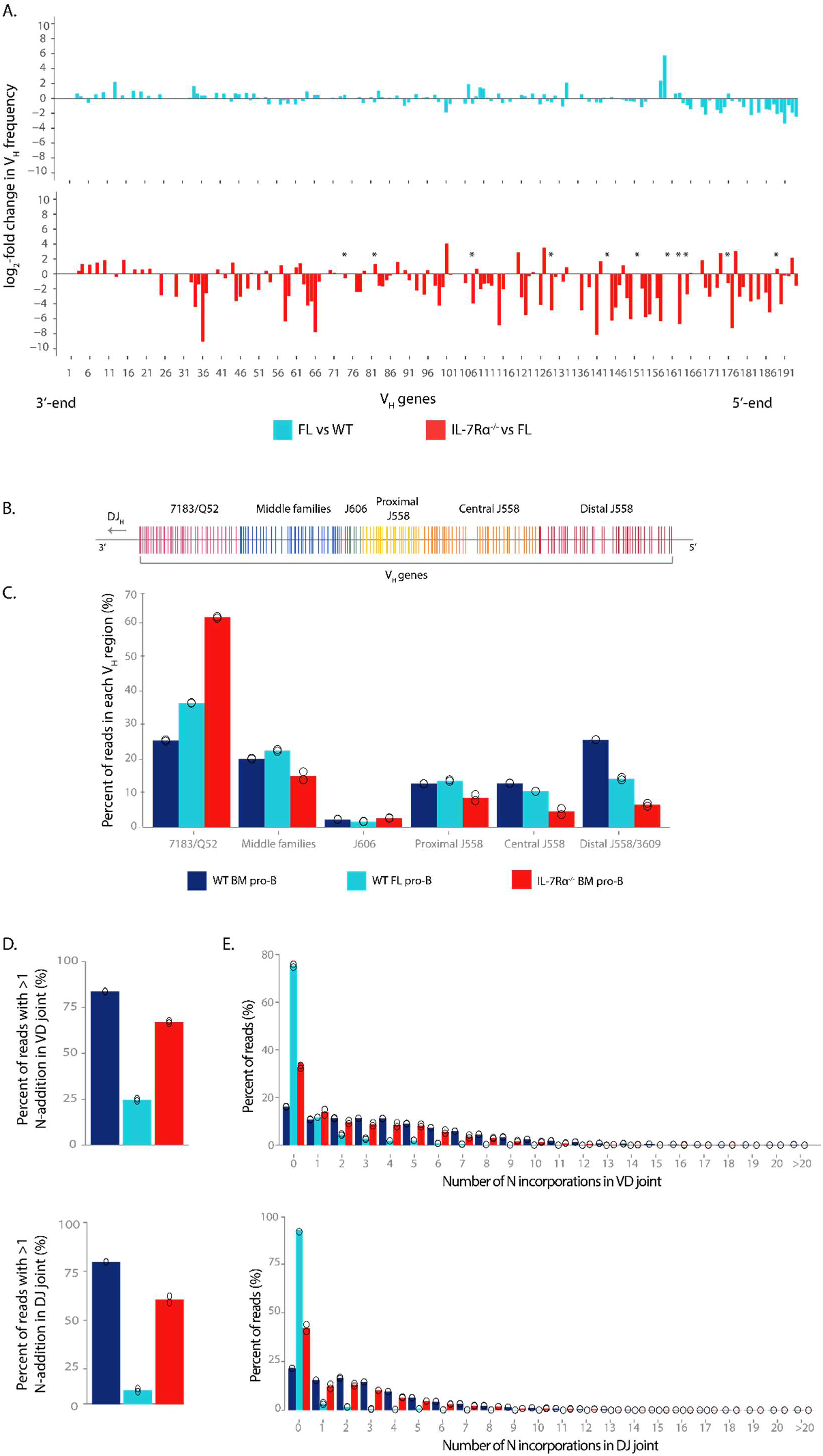
V_H_ recombination frequency in FL pro-B cells is less restricted than IL-7Rα^-/-^ and VDJ sequences show similar levels of N-incorporations greater than FL sequences. A. The average of two WT, two FL and two IL-7Rα^-/-^ replicates was calculated for each V_H_ gene. To display changes between WT and FL frequencies, V_H_ frequencies for FL were divided with the WT mean value and log2-transformed for comparison between models (light blue). Only genes that were active in either genotype are shown. The same was done to compare FL and IL-7Rα^-/-^ frequencies (red). * represents V_H_ genes that had value 0 (only occurring in IL-7Rα^-/-^ replicates). For list of V_H_ gene numbers see supplementary table 1. B. Representation of all V_H_ gene segments as distributed over the V_H_ region (D_H_, J_H_ and C_H_ genes on the left-hand side), coloured by family domains. C. Mapped and quantified VDJ-seq reads over each V_H_ gene were added together for each family domain, and calculated as a percent of total quantified reads for WT (dark blue) and IL-7Rα^-/-^ (red) BM pro-B cells, and wild-type foetal liver (FL – light blue) pro-B cells. Each open circle represents a replicate (2 replicates per genotype). D-E. VDJ-seq libraries were analysed using the webtool IMGT to determine the number of nucleotides inserted into the gene segment junctions during VDJ recombination. Junction between D. V_H_ and D_H_ and E. D_H_ and J_H_ gene segments of WT (dark blue) and IL-7Rα^-/-^ (red) BM pro-B cell recombination events and wild-type FL pro-B cell recombination events (light blue). The number of sequences with more than 1 N-addition (left) and the distribution of sequences with N-additions (right) is shown as a percent of all mapped VDJ-seq sequences. Open circles represent each replicate (2 replicates per genotype).

If we consider V_H_ usage within Igh V_H_ gene family sub-domains (Figure 2B), this is distributed evenly across the locus in WT but is somewhat biased towards the D-proximal 3’ gene families in FL B cells. However, this shift is relatively mild for all but the central and distal J558 genes. Thus, V gene usage appears biased in favour of the 3’ half of the V region in FL B cells (Figure 2C). The IL-7Rα^-/-^ repertoire is also biased towards the 7183/Q52 V_H_ family, but much more so than that of FL B cells. In contrast to FL, this increase in 7183/Q52 V_H_ gene usage was mirrored by a decrease in usage for every other family except the small J606 family. These results demonstrate that recombination events in IL-7Rα^-/-^ cells are markedly more biased towards the 3’ V_H_ genes than FL, suggesting IL-7Rα^-/-^ BM pro-B cells do not simply originate from FL precursors.

To distinguish unambiguously whether VDJ_H_ sequences in IL-7Rα^-/-^ BM cells are derived from FL or adult BM, we analysed the VDJ-seq libraries using IMGT, to compare non-templated (N-) incorporations within VD_H_ and DJ_H_ junctions. Terminal deoxynucleotide transferase (TdT), which is responsible for the insertion of N-nucleotides, is absent in FL and first expressed in pro-B cells in the BM (Feeney et al., 1990; Li et al., 1993). Consistent with this, we identified very few N-additions in FL junctions, with only 25% and 15% of sequences showing more than one incorporation in the VD and DJ junction, respectively, while around 80% of VD and DJ junctions in WT BM had N-additions (Figure 2D). Sequences from IL-7Rα^-/-^ cells also had substantially more N-additions than FL sequences, with around 60% of recombination events showing N-incorporations in both VD and DJ joins. IL-7Rα^-/-^ sequences also showed a similar distribution to WT sequences, including up to 10 additions (Figure 2E). Together these data demonstrate that IL-7Rα^-/-^ pro-B cells undergo V(D)J recombination *de novo* in the BM, but that loss of the IL-7Rα severely restricts usage of V_H_ and D_H_ genes in the formation of the primary Igh repertoire in BM.

### IL-7R signalling does not influence local V gene chromatin state

We next investigated mechanisms by which IL-7R signalling may regulate V(D)J recombination. The reduction in recombination of 5’ V_H_ genes in IL-7Rα^-/-^ BM pro-B cells, together with normal DJ/VDJ ratios, suggests that signalling through IL-7R is specifically required to enable 5’ V_H_ gene recombination, rather than more generally for V(D)J recombination. We first investigated whether altered RSS accessibility could account for the reduced recombination of those V_H_ genes in IL-7Rα^-/-^ cells, since the local enhancer chromatin state is predominantly associated with 5’ V_H_ genes (Bolland et al 2016). We performed ATAC-seq to identify accessible DNA in a chromatin context. We performed these experiments in Rag recombinase-deficient models, which allow the study of pro-B cells that cannot recombine Ig loci, thereby enabling analysis of the intact Igh locus poised for recombination, and avoiding interference from loss of sequence, or elevated V_H_ gene expression, due to recombination. We compared Rag2^-/-^ pro-B cells (which express the endogenous IL-7R) with those from IL-7Rα^-/-^ x Rag2^-/-^ (referred to as IL-7Rα/Rag2^-/-^) mouse BM. Duplicate libraries for both genotypes were highly correlated indicating these data are highly reproducible (Figure S5).

In Rag2^-/-^ pro-B cells, V_H_ RSSs coincided with a peak of accessibility, while the surrounding area was less accessible (Figure 3A). V_H_ RSSs in IL-7Rα/Rag2^-/-^ cells had a similar highly accessible profile, suggesting IL-7R signalling does not activate local accessibility over V_H_ RSSs. The Igκ light chain V gene (Vκ) RSSs were used as a negative control, since the Igκ locus does not become fully activated until the next developmental stage (pre-B) (Matheson et al., 2017). Accordingly, Vκ RSSs were less accessible than V_H_ RSSs (Figure 3A and 3B). Notably, the pattern for Igκ Vκ genes was similar in Rag2^-/-^ and IL-7Rα/Rag2^-/-^ pro-B cells, suggesting that these are not activated prematurely in the absence of the IL-7Rα.

**Figure 3:**
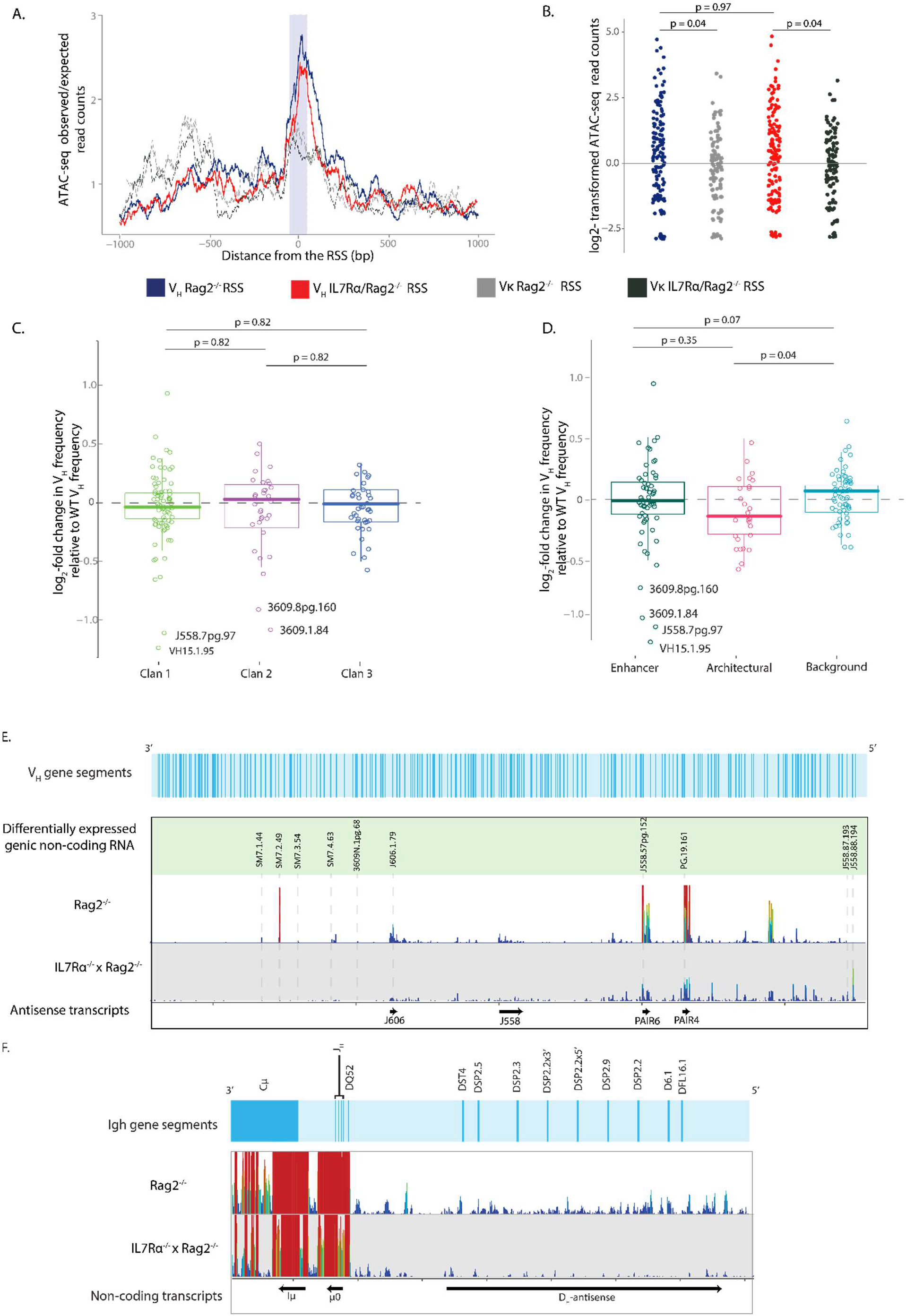
IL-7Rα^-/-^Rag2^-/-^ pro-B cell V_H_ RSSs show no significant changes in RSS accessibility relative to Rag2^-/-^ V_H_ RSSs but show differences in non-coding transcription over the Igh locus in Rag2^-/-^ and IL-7Rα^-/-^Rag2^-/-^ pro-B cells. A. Profile of accessibility over the V_H_ and Vκ RSSs in Rag2^-/-^ and IL-7Rα^-/-^Rag2^-/-^ pro-B cells. Accessibility tracks are shown over a 1000 bp region centred on the RSS. ATAC-seq reads were quantified over each bp and each track is an average for all Rag2^-/-^ V_H_ (blue) and Vκ (dotted grey) RSSs, and IL-7Rα^-/-^/Rag2^-/-^ V_H_ RSSs (red) and Vκ (dotted black). Average counts per bp were calculated relative to the total average counts over the whole region (2000 bp). Purple area represents the location of the RSS. B. ATAC-seq reads over a 50 bp region centred over V_H_ (n = 195) and Vκ (n = 162) RSS for Rag2^-/-^ and IL-7Rα/Rag2^-/-^ were quantified and log2 transformed using DESeq2. ANOVA (degrees freedom = 3, F-value = 3.68, p-value = 0.012). The differential expression values for each RSS as calculated by DESeq2 were grouped by C. evolutionary origin: clan 1 (n = 78), clan 2 (n = 27) and clan 3 (n = 26), ANOVA (degrees freedom = 2, F-value = 0.27, p-value = 0.77); D. chromatin state: enhancer (n = 68), architectural (n = 30) and background (n = 33); Kruskal-Wallis test (degrees freedom = 2, chi-squared = 7.51, p-value = 0.023) followed by pairwise Wilcox test (adjusted by Benjamini and Hochberg method) to calculate p-value. E-F. RNA-seq reads for Rag2^-/-^ and IL-7Rα^-/-^Rag2^-/-^ were quantified per 60 bp bins along the Igh locus and normalised by reads per million (generated using Seqmonk). The height and colour of each bar represents a relative number of reads over each probe: high red bars have more reads than short blue bars, with ranging colours and heights representing intermediate quantities of reads. Each track was generated from an average of two RNA-seq libraries. E. Transcription over the V_H_ region of the Igh. The top of the figure shows the location of all V_H_ gene segments (light blue track). Transcription over ten V_H_ genes which were significantly differentially expressed (see Table 2) is shown in the green track (grey dotted line marking their location), and the location of antisense intragenic non-coding transcripts are shown as black arrows. F. Transcription over the D_H_, J_H_ and C_H_ regions of the Igh. D_H_, J_H_ and C_H_ gene segment locations are shown in the light blue track. Intergenic non-coding transcripts are shown as black arrows at the bottom the figure.

We then divided the RSSs into clans and chromatin states to test if RSS from these groups showed different accessibility patterns in IL-7Rα/Rag2^-/-^. This analysis showed no significant difference between clans (Figure 3C). RSS accessibility in the enhancer and architectural groups was significantly reduced compared to the background state V_H_ genes, which showed a small increase in RSS accessibility relative to WT (Figure 3D). Taken together, these data indicate little difference in local accessibility at V_H_ genes in the absence of the IL-7Rα.

### Igh antisense intergenic transcription, but not V_H_ genic transcription, is impaired in IL-7Rα^-/-^ pro-B cells

Because of the precise developmental timing of its appearance and disappearance, non-coding transcription has been proposed to promote chromatin accessibility to facilitate Igh V gene recombination (Yancopoulos & Alt 1985, Corcoran et al., 1998; Bolland et al., 2004). The Igh locus has two types of non-coding transcripts in the V_H_ region: short genic sense transcripts which begin at V_H_ promoters (Yancopoulos & Alt 1985; Corcoran et al. 1998), and long intergenic antisense transcripts (Bolland et al. 2004; Verma-Gaur et al. 2012; Bolland et al. 2016). To investigate changes in non-coding transcription in the absence of IL-7R signalling, we performed RNA-seq on Rag2^-/-^ and IL-7Rα/Rag2^-/-^ pro-B cells (Figure S6a-c). This revealed that there is generally little V_H_ genic transcription over the 3’ V genes with 31 of the 39 most D-proximal V genes showing no transcription in Rag2^-/-^ pro-B cells. Very few non-coding V_H_ genic transcripts were differentially expressed between IL-7Rα/Rag2^-/-^ and Rag2^-/-^ cells (Table 1 - for full results see S Table 2). Notable exceptions included a few V_H_ genes in the middle-families domain (including all four members of the SM7 family), which were highly abundant in Rag2^-/-^, but almost completely absent in IL-7Rα/Rag2^-/-^ cells. Conversely, two of the most 5’ V_H_ genes, J558.88.194 and J558.87.193, which are not transcribed in Rag2^-/-^ pro-B cells, are highly expressed in IL-7Rα/Rag2^-/-^ pro-B cells (Figure 3E).

**Table 1:** Significantly differentially expressed transcripts over V_H_ gene segments in Rag2^-/-^ and IL-7Rα^-/-^Rag2^-/-^ pro-B cells. RNA-seq raw reads were quantified over each V_H_ gene probe. Each of two replicates was quantified separately to determine differences in transcription between Rag2^-/-^ and IL-7Rα^-/-^ Rag2^-/-^ pro-B cells. The adjusted mean for each probe, as well as the log2-fold change and adjusted p-value in the IL-7Rα^-/-^ Rag2^-/-^ relative to Rag2^-/-^ samples was calculated by DESeq2. See supplementary Table 2 for full results of analysis.

| V<sub>H</sub> gene mean | Rag2<sup>-/-</sup> mean | IL-7Rα/Rag2<sup>-/-</sup> Fold Change | log2 | p-adj |
| --- | --- | --- | --- | --- |
| J558.88.194 | 0 | 104.62 | 5.94 | 9.29E-06 |
| J558.87.193 | 2.24 | 53.11 | 3.32 | 2.97E-02 |
| PG.19.161 | 308.57 | 26.41 | -3.37 | 9.78E-08 |
| J558.57pg.152 | 464.11 | 34.51 | -3.64 | 8.52E-15 |
| J606.1.79 | 127.48 | 9.12 | -3.55 | 1.69E-06 |
| 3609N.1pg.68 | 17.78 | 0 | -3.84 | 2.31E-02 |
| SM7.4.63 | 19.47 | 0.44 | -3.38 | 4.34E-02 |
| SM7.3.54 | 17.68 | 0 | -3.86 | 2.15E-02 |
| SM7.2.49 | 362.92 | 5.8 | -5.58 | 1.85E-16 |
| SM7.1.44 | 57.96 | 2.19 | -3.8 | 2.58E-03 |

Long intergenic antisense non-coding transcripts in the V_H_ (PAIR4, PAIR6, J558 and J606) and DJ_H_ regions (Iμ, μ0 and D_H_-antisense) were also analysed. Although the RNA-seq libraries generated were not strand-specific, these known transcripts are easily distinguished from the much less frequently transcribed genic transcripts in Rag2^-/-^ pro-B cells (Figure 3E). Strikingly, antisense transcription throughout the V_H_ region was almost completely absent in IL-7Rα/Rag2^-/-^ pro-B cells, and in some cases was difficult to distinguish from the surrounding genic transcription. DESeq2 differential expression analysis revealed a significant reduction in all V_H_ antisense transcripts tested (Table 2). Furthermore, D_H_ antisense transcription was almost completely lost over the entire DJ_H_ region (Figure 3F), while sense transcription over the J_H_ (μ0 transcript) or the C_H_ (Iμ transcript) regions was relatively unchanged in IL-7Rα/Rag2^-/-^ relative to Rag2^-/-^ cells (Table 2).

**Table 2:** Significantly differentially expressed transcripts over Igh V intergenic regions in Rag2^-/-^ and IL-7Rα^-/-^Rag2^-/-^ pro-B cells. RNA-seq raw reads were quantified over intergenic non-coding transcript probes (see Figure 3E – length of probes shown in bp). Each of two replicates was quantified separately to determine differences in transcription between Rag2^-/-^ and IL-7Rα^-/-^ Rag2^-/-^ pro-B cells. The adjusted mean for each probe, as well as the log2-fold change and adjusted p-value in the IL-7Rα^-/-^Rag2^-/-^ relative to Rag2^-/-^ samples was calculated by DESeq2. Transcripts that were significantly differentially expressed are marked with *.

| Intergenic transcript | Probe length (bp) | Rag2<sup>-/-</sup> mean | IL-7Rα/Rag2<sup>-/-</sup> mean | log2 Fold Change | p-adj |
| --- | --- | --- | --- | --- | --- |
| RNA-PAIR4* | 26,160 | 8408.81 | 1140.45 | -2.79 | 3.23E-08 |
| RNA-PAIR6* | 23,314 | 18185.29 | 1951.23 | -3.1 | 3.17E-08 |
| RNA-J558* | 89,679 | 1043.85 | 186.02 | -2.44 | 1.95E-09 |
| RNA-J606* | 29,461 | 1475.66 | 213.42 | -2.74 | 2.49E-13 |
| RNA-D<sub>H</sub> region* | 57,688 | 1149.29 | 248.44 | -2.09 | 5.17E-03 |
| RNA - μ0 | 2,572 | 8381.6 | 4372.01 | -0.91 | 1.62E-01 |
| RNA-Iμ | 4,152 | 55090.86 | 28612.49 | -0.91 | 2.80E-01 |

### EBF1, PAX5 and other key B cell lineage-specifying genes are mis-regulated in IL-7Rα^-/-^ pro-B cells

We next examined genome-wide alterations in gene expression in the absence of the IL-7R. Out of 4,793 differentially expressed genes in IL-7Rα/Rag2-/- pro-B compared with Rag2-/- pro-B cells, 3,780 genes were upregulated, while 1,013 genes were downregulated (Figure S6d). EBF1 and PAX5 are two of the most important transcription factors driving B cell specification and commitment (Pongubala et al., 2008; Fuxa et al., 2007; Rumfelt et al., 2006; O’Riordan, & Grosschedl., 1999) and both have been implicated as effectors of IL-7R signalling (Kikuchi et al., 2005; Roessler et al., 2007; Decker et al., 2009; Corcoran et al., 1998). Consistent with previous reports, expression of both *Ebf1* and *Pax5* was substantially reduced in IL-7Rα/Rag2^-/-^ relative to Rag2^-/-^ pro-B cells (Table 3). Importantly, several key B-lineage genes regulated by *Ebf1* and *Pax5* had reduced transcription levels in IL-7Rα/Rag2^-/-^ cells, including *Foxo1, Rag1* and *Cd79a* (which codes for Igα, part of the pre-B cell receptor complex), previously shown to be downregulated in EBF1-deficient cells (Györy et al., 2012). PAX5-activated genes including *Smarca4* (which encodes BRG1) and *Lef1* were also decreased. Conversely, FLT3R (*Flt3*), active in pre-pro-B cells and downregulated in pro-B cells in a PAX5-dependent manner (Pridans et al., 2008), was upregulated in IL-7Rα/Rag2^-/-^ pro-B cells.

**Table 3:** EBF1, PAX5 and their key targets are differentially expressed in IL-7Rα^-/-^ Rag2^-/-^ relative to Rag2^-/-^ pro-B cells. RNA-seq raw reads were quantified over all genes. Each of two replicates was quantified separately and analysed to determine differences in transcription between Rag2^-/-^ and IL-7Rα^-/-^ Rag2^-/-^ pro-B cells. The adjusted mean for each probe, as well as the log2-fold change and adjusted p-value in the IL-7Rα^-/-^ Rag2^-/-^ relative to Rag2^-/-^ samples was calculated by DESeq2.

| Gene | Rag2<sup>-/-</sup> mean | IL-7Rα/Rag2<sup>-/-</sup> mean | log2 FoldChange | p-adj |
| --- | --- | --- | --- | --- |
| Ebf1 | 198467.7 | 31632.21 | -2.57 | 2.79E-07 |
| Pax5 | 22013.66 | 5049.33 | -2.05 | 5.39E-04 |
| Flt3 | 371.14 | 2680.54 | 2.8 | 1.31E-14 |
| Irf8 | 1727.72 | 7817.74 | 2.15 | 1.34E-09 |
| Ikzf3 | 453.74 | 2896.93 | 2.58 | 6.89E-06 |
| Lef1 | 50047.29 | 7152.81 | -2.69 | 1.39E-05 |
| Foxo1 | 19360.34 | 5176.08 | -1.86 | 2.37E-05 |
| Cd79a | 36517.38 | 5427.5 | -2.6 | 2.64E-04 |
| Rag1 | 24672.56 | 4710.95 | -2.26 | 2.37E-03 |
| Smarca4 | 94899.66 | 25252.66 | -1.84 | 3.19E-03 |
| Ezh2 | 21004.9 | 8795.46 | -1.23 | 3.77E-03 |
| Myb | 29838.38 | 16868.3 | -0.8 | 2.41E-01 |
| Irf4 | 5144.04 | 2515.98 | -0.98 | 3.14E-01 |
| Pdcd1 | 0 | 2.37 | 0.86 | 7.68E-01 |

However, other B cell-specific genes that are under PAX5 and EBF1 control were expressed normally: *Irf4* (a direct target of PAX5 and EBF1), *Myb* (reduced in EBF1-deficient cells) and *Pdcd1* (repressed by EBF1) showed no significant transcriptional changes between IL-7Rα/Rag2^-/-^ and Rag2^-/-^ cells (Györy et al., 2012; Pridans et al., 2008; Treiber et al., 2010). Indeed, *Irf8* and *Ikzf3* (Aiolos), which are also PAX5-activated targets and are directly bound by EBF1, were both more highly expressed in IL-7Rα/Rag2^-/-^ cells. Together, these results suggest that EBF1 and PAX5 function is mis-regulated in IL-7Rα^-/-^ cells. To determine whether IL-7R signalling additionally influences the binding pattern of these and other TFs, we compared accessible hypersensitivity sites between the genotypes with ATAC-seq, using the MACS caller (Zhang et al., 2008) function in Seqmonk within our ATAC-seq datasets. We divided the sites into two groups: those which had fewer reads (less accessible), and those which had more reads (more accessible) in IL-7Rα/Rag2^-/-^ than Rag2^-/-^ cells. We then used Hypergeometric Optimization of Motif EnRichment (HOMER) to identify TF motifs within these sites. This allowed us to infer TFs which are binding less often (sites which are less accessible) in IL-7Rα/Rag2^-/-^relative to Rag2^-/-^, and vice versa. This identified EBF1 as binding less often in IL-7Rα/Rag2^-/-^ cells relative to Rag2^-/-^ cells, correlating with a reduction in its expression and in the expression of EBF1-target genes. The motif for PAX5 was not found in this analysis, although the PAX8 motif (which has a similar binding pattern to PAX5) was detected. We infer that this is in fact PAX5, since PAX8 is not expressed in B cells. Again, our analysis found reduced representation of this motif in ATACseq-accessible sites in 7Rα/Rag2^-/-^ cells (Table 4A). This analysis also detected reduced accessibility at motifs of other important TFs involved in B cell specification and development, including E2A, which activates many B-lineage genes and represses non-B-lineage genes (Dias et al., 2008; Kee et al., 1998; Ikawa et al., 2004; Lin et al., 2010).

**Table 4:**
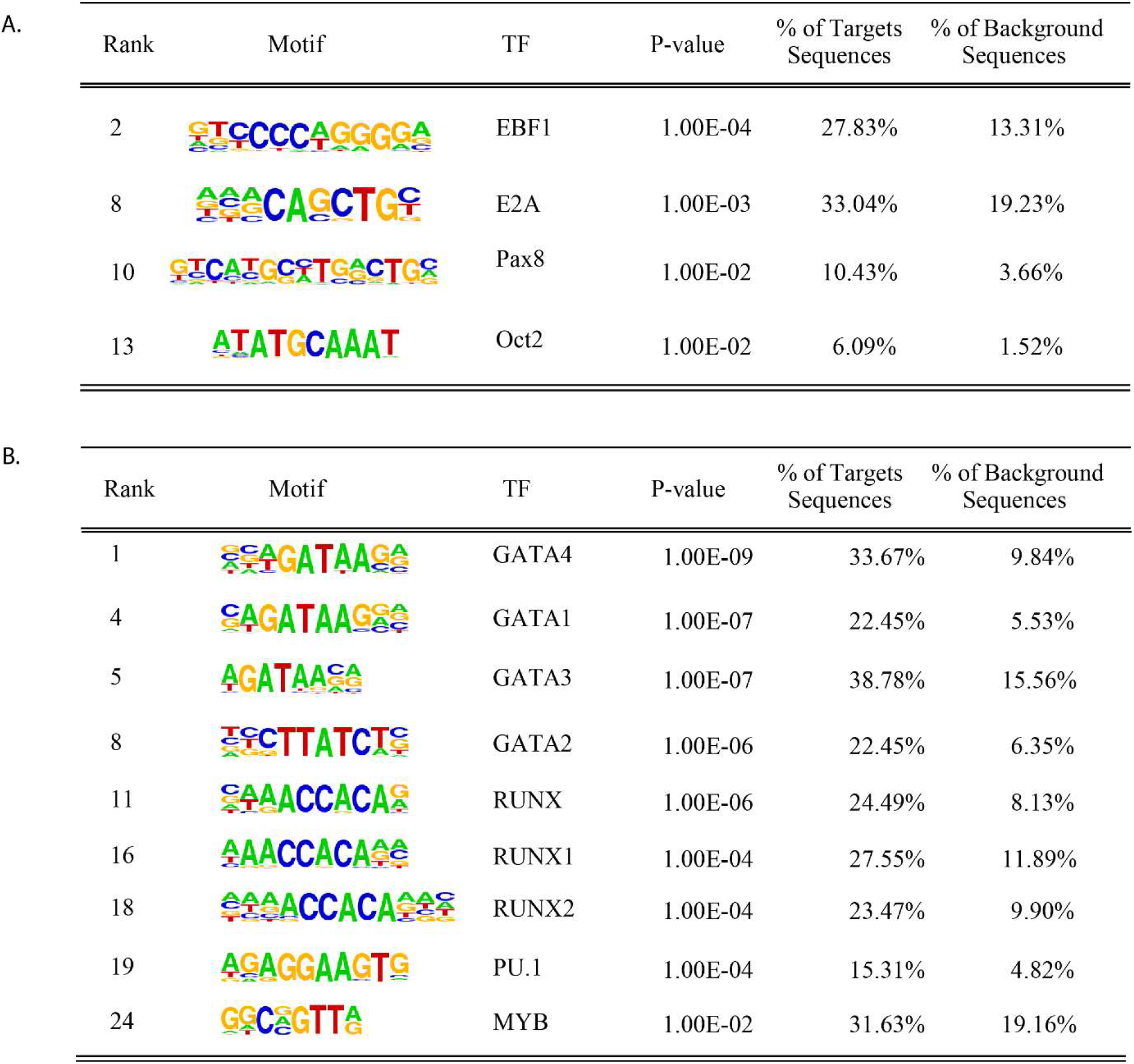
TF motif analysis at genomic sites of altered accessibility. Peaks of accessibility were identified from ATAC-seq data by the MACS peak calling function in Seqmonk. **A.** Sites that were less accessible in IL-7Rα/Rag2^-/-^ relative to Rag2^-/-^ cells were analysed using DESeq2 and TF motif enrichment was carried out using HOMER. Relevant significantly enriched motifs are shown in order of significance (rank indicated their position in the list), with the motif, TF name, P-value, percent of target regions which contained this motif, and percent of background regions which had this motif. **B.** The same analysis was done as in A for peaks that were more accessible in IL-7Rα/Rag2^-/-^ relative to Rag2^-/-^ cells.

|  |  |  |  |  |  |  |
| --- | --- | --- | --- | --- | --- | --- |
| A. | Rank | Motif | TF | P-value | % of Targets Sequences | % of Background Sequences |
|  | 2 |  | EBF1 | 1.00E-04 | 27.83% | 13.31% |
|  | 8 |  | E2A | 1.00E-03 | 33.04% | 19.23% |
|  | 10 |  | Pax8 | 1.00E-02 | 10.43% | 3.66% |
|  | 13 |  | Oct2 | 1.00E-02 | 6.09% | 1.52% |
| B. | Rank | Motif | TF | P-value | % of Targets Sequences | % of Background Sequences |
|  | 1 |  | GATA4 | 1.00E-09 | 33.67% | 9.84% |
|  | 4 |  | GATA1 | 1.00E-07 | 22.45% | 5.53% |
|  | 5 |  | GATA3 | 1.00E-07 | 38.78% | 15.56% |
|  | 8 |  | GATA2 | 1.00E-06 | 22.45% | 6.35% |
|  | 11 |  | RUNX | 1.00E-06 | 24.49% | 8.13% |
|  | 16 |  | RUNX1 | 1.00E-04 | 27.55% | 11.89% |
|  | 18 |  | RUNX2 | 1.00E-04 | 23.47% | 9.90% |
|  | 19 |  | PU.1 | 1.00E-04 | 15.31% | 4.82% |
|  | 24 |  | MYB | 1.00E-02 | 31.63% | 19.16% |

This reduction in B cell specification was also reflected in TF motifs within sites which are more accessible in IL-7Rα/Rag2^-/-^ cells relative to Rag2^-/-^ pro-B cells. T cell development programme TF motifs, including several GATA family TFs, as well as early B cell priming TFs including PU.1, MYB and RUNX, were associated with more accessible regions in IL-7Rα/Rag2^-/-^ pro-B cells (Table 4B). Overall, the pattern of TF motif accessibility suggests that a function of the IL-7Rα in pro-B cells is to enforce commitment to the B cell lineage.

## Discussion

In this study, against a background of conflicting reports in the literature, we set out to determine whether and how the IL-7R plays a role in *de novo* immunoglobulin gene recombination in BM B cells. We compared the Igh repertoire in BM pro-B cells lacking the signalling portion of the IL-7R (IL-7Rα^-/-^) with the repertoire in WT BM and FL pro-B cells. As previously reported (Jeong and Teale., 1988; Yancopoulos et al., 1988), we found that pro-B cells from FL cells showed a bias towards usage of V genes from the 3’-end of the V_H_ region. However, this bias was much less than expected, since virtually all recombining V genes in WT BM also participated in recombination in FL, albeit in reduced proportions in some cases. Thus, we show for the first time that the FL B cell antibody repertoire is far less restricted than previously thought. Our study calls into question the current model of B cell ontogeny, in which complex antibody repertoires do not develop until a few weeks of age in the mouse, or a few months in human (Siegrist and Aspinall 2009). It will be important to investigate which mechanisms that underpin adult Igh repertoire formation are already in place in FL, including long-range 3D looping of the V region, and local V gene activation. These mechanisms are dependent on factors including CTCF, PAX5 and YY1 in BM B cells (Gerasimova et al., 2015; Bolland et al., 2016; Jain et al., 2018). Since PAX5 is essential for FL B cell development and Igh V_H_ to DJ_H_ recombination (Nutt et al., 1997), it may play similar roles in local V gene activation and/or long-range looping in FL B cells.

The unprecedented depth of analysis of individual IL-7Rα^-/-^ VDJ_H_ and DJ_H_ sequences we have achieved here by means of VDJ-seq demonstrates that IL-7Rα^-/-^ pro-B cells display widespread defects at both stages of Igh recombination. Most importantly, the V_H_ repertoire was highly biased towards 3’ V_H_ genes, to the detriment of both middle and 5’ V_H_ genes. Reduced use of 5’ V_H_ genes was much more pronounced in IL-7Rα^-/-^ BM pro-B cells than in WT FL pro-B cells, indicating that IL-7R signalling is specifically needed in the BM to make all V genes available to the primary antibody repertoire. The presence of N-additions within the VDJ_H_ and DJ_H_ sequences obtained from IL-7Rα^-/-^ cells is an important demonstration that they are derived from BM rather than FL progenitors. Thus, our findings concur with a previous study in neonatal IL-7Rα^-/-^ BM (Hesslein et al., 2006), and disagree with reports that suggested a complete block in B cell development in IL-7Rα^-/-^ BM (Kikuchi et al., 2005; Peschon et al., 1994; Miller et al., 2002; Carvalho et al., 2001). Because the IL-7R is required for proliferation and survival, the latter studies inferred that B cells progressing through development had originated in the FL, where the IL-7R is not essential (Erlandsson et al., 2004). We agree that loss of the IL-7R severely depletes BM B cell numbers, but we show here that V(D)J recombination continues to occur, enabling us to uncover very specific roles of the IL-7R in D_H_ and V_H_ gene usage in the Igh repertoire.

It is noteworthy that our data show N-additions were also detected at the D_H_-J_H_ junction, indicating that TdT is expressed from the CLP stage in IL-7Rα^-/-^ BM, and that D_H_ to J_H_ rearrangements also took place *de novo* in the BM. Thus, in the absence of IL-7Rα, V(D)J recombination progressed with normal dynamics but with severely restricted participation of V_H_ and D_H_ genes. A higher percentage of IL-7Rα^-/-^ sequences (20%) had no N-additions relative to WT, but this is likely to be due to the difference in age between the WT (12 weeks) and IL-7Rα^-/-^ (6-8 weeks) mice analysed, since recombination junctions in the BM are less complex early after birth, and progressively become more diverse with age (Delassus et al., 1998). Since most sequences from IL-7Rα^-/-^ cells have the properties of WT BM-derived B cell sequences, we conclude that the impaired V(D)J repertoire observed in IL-7Rα^-/-^ BM B cells is specifically due to loss of stage-specific IL-7R function in the BM.

Our results are also at odds with the conclusions of Malin et al., (2010), who suggested that IL-7R signalling is not required for recombination in BM B cells. It is important to note, however, that their study focussed on the role of STAT5, and did not account for PI3K or other signalling mechanisms engaged by the IL-7R. Moreover, RAG1-Cre deletion of *Stat5* was not complete in CLPs which may have expressed sufficient STAT5 to establish the necessary positive feedback loop between EBF1 and PAX5 (Zandi et al., 2010). The authors showed that IL-7Rα^-/-^ B cell development could be partially rescued using a vav-cre expressed bcl2 transgene, suggesting a crucial role for the IL-7R in survival of CLPs. However, *Bcl2* under the control of the Igh Eμ intronic enhancer did not rescue B cell development (Maraskovsky et al., 1998). Thus, the IL-7R has important functions beyond survival at the stage at which V(D)J recombination is taking place. Furthermore, the IL-7R does not require STAT5 activation or PI3K signalling to promote V(D)J recombination *in vitro* (Corcoran et al., 1996).

Our study has uncovered mechanisms underpinning impaired recombination of 5’ and middle V_H_ genes. Local accessibility of antigen receptor gene segments for recombination is thought to be influenced by several mechanisms including nucleosome remodelling (Bevington and Boyes., 2013; Pulivarthy et al., 2016), which in turn could modulate or be modulated by the activity of factors associated with local V gene active chromatin states (Bolland et al., 2016). Here, ATAC-seq showed that while there was a wide range of accessibility values among V_H_ genes within different clans and chromatin states in both genotypes, there were no significant differences between clans or states in either genotype, and overall local accessibility over V_H_ gene RSSs was unchanged in IL-7Rα/Rag2^-/-^ relative to Rag2^-/-^ cells. In both models there was very low accessibility at Vκ gene RSSs, consistent with Vκ genes being inactive until the next developmental stage. Importantly, this suggests that the surviving IL-7Rα^-/-^ B cells have not ‘rushed through’ to the IL-7R-independent pre-B cell stage where increased Vκ access would be expected. Interestingly, it has been shown that IL-7R signalling must be downregulated to enable Igκ accessibility and recombination (Johnson et al., 2008; Mandal et al., 2011). In our hands, the lack of increased accessibility at Vκ gene RSSs in IL-7Rα/Rag2^-/-^ B cells indicates that loss of the IL-7R is not sufficient to activate Vκ genes, and suggests that additional mechanisms must be at play, perhaps involving CXCR4, recently shown to activate Igκ (Mandal et al., 2019). Overall, we find no evidence to indicate that defects in local accessibility at the 5’ and middle V genes account for the preference for recombination of 3’ V_H_ genes in IL-7Rα^-/-^ cells.

Our detailed analysis of Igh V_H_ genic sense non-coding transcription in IL-7Rα^-/-^ pro-B cells revealed that this transcription generally did not change. The few changes observed did not correlate with the skewed recombination found in the absence of IL-7R signalling. This corroborates recent integrated multiomics studies that found no correlation between V_H_ genic non-coding transcription and V_H_ usage frequency in normal Igh recombination (Choi et al., 2013; Bolland et al., 2016). Earlier studies using PCR-based methods (Corcoran et al., 1998; Bertolino et al., 2005) showed reduced sense transcription over 5’ V_H_ genes in IL-7Rα^-/-^ cells. These studies used primers that did not distinguish between sense and then undiscovered antisense intergenic transcripts in the Igh locus. Since the latter overlap 5’ V_H_ gene segments (Bolland et al., 2004), and are transcribed at levels orders of magnitude higher than genic transcripts (Choi et al., 2013; Bolland et al., 2016), the previous reports probably identified the downregulation of these antisense transcripts. Indeed, several of the few V genes observed here with reduced sense transcripts overlap regions of antisense transcription.

Non-coding intergenic transcription has been shown to activate recombination of corresponding Jα genes in the TCRα locus *in vivo* (Abarrategui and Krangel, 2006), while *de novo* antisense transcription over 3’ V_H_ genes increases Igh 3’ V_H_ gene recombination (Guo et al., 2011). Together, these and other findings have supported a model in which V intergenic transcription drives recombination (Corcoran et al., 2010). Here we found widespread loss of all of the major PAX5-dependent (PAIRs 4 and 6) and PAX5-independent (J558, J606) antisense intergenic transcripts (Bolland et al., 2004; Ebert et al., 2011; Verma-Gaur et al., 2012; Choi et al., 2013; Bolland et al., 2016). This suggests that the IL-7R regulates expression of all Igh antisense transcripts and downstream functions, and supports a role for the receptor in promoting long-range mechanisms underpinning V_H_ to D_H_ recombination. We did not observe *de novo* antisense transcription over 3’ V_H_ genes in IL-7Rα^-/-^ pro-B cells. Importantly, this is consistent with the relative increase in 3’ V_H_ gene recombination being secondary to the defect in 5’ recombination, rather than a *bona fide* increase in 3’ recombination (Guo et al., 2011).

IL-7R signalling, mediated by Pax5, may also influence V(D)J recombination through contraction and looping of the Igh locus. PAX5, downregulated in IL-7Rα^-/-^ cells, is essential for Igh locus contraction and looping (Fuxa et al., 2004; Nutt et al., 1997; Hesslein et al., 2003; Medvedovic et al., 2013; Montefiori et al., 2016). PAX5 has pleiotropic functions, but the specific downregulation of PAX5-dependent PAIR transcription observed here in the IL-7Rα^-/-^ suggests that PAX5 binding and function at these regulatory regions is directly impaired (Ebert et al., 2011).

A key insight provided by VDJ-seq was that V(D)J recombination in IL-7Rα^-/-^ cells is compromised as early as the pre-pro-B stage, when D_H_ to J_H_ recombination is taking place. Representation of several of the non-flanking DSP family of D_H_ genes was severely reduced on DJ recombined alleles in IL-7Rα^-/-^ pro-B cells. This finding is at odds with previous analyses, but these did not have sufficient resolution to distinguish between DSP gene family members (Corcoran et al., 1998; Bertolino et al., 2005). Strikingly, antisense non-coding transcription over the D_H_ region was also severely reduced. Conversely, recombination of the most 3’ D gene, DQ52, located in a separate chromatin domain shared with the J_H_ genes, where transcription was unaffected by loss of the IL-7R, was proportionally increased. These findings support our previous model that antisense transcription over the DSP genes activates their recombination (Bolland et al., 2007). They provide additional evidence against the opposing model that this transcription represses recombination of the DSPs (Chakraborty et al., 2007), and reveal a new role for the IL-7R in activating D_H_ antisense transcription to drive D_H_ to J_H_ recombination.

Our genome-wide RNA-seq and ATAC-seq analyses also allowed us to look in detail at key effectors of IL-7R signalling, and their targets. We confirmed that the IL-7R regulates EBF1 and PAX5, and provided a detailed picture of their targets that are dysregulated by loss of IL-7R signalling. Importantly, analysis of TF binding motifs in regions of increased and decreased accessibility, albeit *in silico*, gave powerful conceptual insights. Reduced accessibility at putative EBF1 binding sites pointed to specific concrete consequences of reduced EBF1 expression. Conversely, increased accessibility at TF motifs associated with T cell development suggests that, although IL-7Rα^-/-^ B cells are committed to the B cell lineage in the sense that they have undergone V to DJ recombination, they nevertheless may remain plastic. This is reminiscent of PAX5^-/-^ B cells which also can revert to other lineages despite undergoing V(D)J recombination (Nutt et al., 1999).

Our findings also have important implications for understanding of human B cell development. Studies of early paediatric severe combined immunodeficiency disease (SCID) patients suggested that, in surprising contrast to mouse, human T cell development required IL-7, but human B cell development did not (LeBien, 2000). However, more recent studies have shown that while fetal and neonatal human B cell development can proceed without IL-7, adult B cell development is critically dependent on IL-7R signalling, thereby closely aligning the dynamics of mouse and human IL-7R dependency (Parrish et al., 2009; Milford et al., 2016). These studies focused on B cell numbers and progression through development, and underlying mechanisms are unclear. It will be important to determine whether the IL-7R plays a role in immunoglobulin gene recombination during human B cell development. Our findings provide an important way forward for investigations in human immunodeficiency diseases and ageing, both characterised by restricted antibody repertoires and poor antibody response to infection (Siegrist & Aspinall 2009; Martin et al 2015). Indeed, both Igh recombination and IL-7R signalling are impaired in ageing in mouse (Stephan et al 1997; Szabo et al 1999) and human, and the role and therapeutic potential of the IL-7R in human ageing is an emerging area of interest (Passtoors et al., 2015). Our findings make a strong case for investigating the potential of IL-7 for boosting antibody repertoires therapeutically, since we show that its receptor underpins the diversity of the naïve antibody repertoire that responds to new infections.

In conclusion, we reveal that IL-7R signalling shapes the Igh repertoire at both the D_H_-to-J_H_ and V_H_-to-DJ_H_ stages of Igh V(D)J recombination in mouse BM and identify several mechanisms by which IL-7R signalling can activate the Igh locus. IL-7R signalling is therefore essential for promoting the use of the wide repertoire of V_H_ and D_H_ gene segments during V(D)J recombination, which expands antibody diversity and ensures a robust activation of the adaptive immune system.

## Methods

### Mice

RAG2^-/-^ (Shinkai et al., 1992), IL-7Rα^-/-^ (Peschon et al., 1994) and IL-7Rα^-/-^ crossed with RAG2^-/-^ (IL-7Rα^-/-^ x RAG2^-/-^) C57BL/6 mice were maintained in accordance with Babraham Institute Animal Welfare and Ethical Review Body and Home Office rules under Project Licence 80/2529. Recommended ARRIVE reporting guidelines were followed. Mice were bred and maintained in the Babraham Institute Biological Services Unit under Specific Opportunistic Pathogen Free (SOPF) conditions. After weaning, mice were maintained in individually ventilated cages (2-5 mice per cage). Mice were fed CRM (P) VP diet (Special Diet Services) ad libitum, and millet, sunflower or poppy seeds at cage-cleaning as environmental enrichment. Health status was monitored closely and any mouse with clinical signs of ill-health or distress persisting for more than three days was culled. Treatment with antibiotics was not permitted to avoid interference with immune function. Thus, all mice remained ‘sub-threshold’ under UK Home Office severity categorization. 6-8-week-old IL-7Rα^-/-^ and IL-7Rα^-/-^ x RAG2^-/-^ mice (all mixed sex), and 10-12 week old RAG2^-/-^ mice (one female replicate and one male replicate) were used. Although wild-type (WT) comparison data was from 12 week old mice, IL-7Rα^-/-^ animals were taken before 10 weeks because they produce fewer BM B cells as they age, with very few produced after 10 weeks (Peschon et al., 1994; Erlandsson et al., 2004). To maximise cell numbers and considering IL-7Rα^-/-^ mice as young as 3 weeks have adult B cell populations (Hesslein et al., 2006), pro-B cells from 6-8-week old mice were taken for sorting. Fetal livers (FL) were harvested from day 15.5 mouse embryos.

### Primary cells

BM was depleted of macrophages, granulocytes, erythroid lineage and T cells using biotinylated antibodies against CD11b (MAC-1; ebioscience), Ly6G (Gr-1; ebioscience), Ly6C (Abd Serotec), Ter119 (ebioscience) and CD3e (ebioscience) followed by streptavidin MACs beads (Miltenyi). MACS depletion was carried out as for BM, using TER119-biotin at a higher concentration (1:200). Thereafter, pro-B cells from IL-7Rα^-/-^ BM and WT FL were flow-sorted as B220^+^ CD19^+^, while IL-7Rα^-/-^ x RAG2^-/-^ and RAG2^-/-^ BM B pro-B cells were sorted as a B220^+^ CD19^+^ CD43^+^ population on a BD FACSAria in the Babraham Institute Flow Cytometry facility. WT BM pro-B cells were sorted as B220^+^ CD19^+^ CD25^-^ sIgM^-^ CD43^+^ (Bolland et al., 2016). Sorting antibodies were from BD Bioscience.

### DNA extraction

Genomic DNA was isolated from mouse B cells using the DNeasy kit (Qiagen) according to the manufacturer’s instructions.

### RNA-seq

Total RNA was extracted from ~200,000 cells for each replicate using the RNeasy Plus kit (Qiagen). cDNA preparation was performed using the Ovation V2 kit (NuGen) protocol, and 200 ng of cDNA carried through to generate 50bp paired-end RNA-seq libraries for Illumina sequencing as described in Parkhomchuk et al., (2009) except polyA+ RNA selection and strand-specificity steps were omitted, and first strand cDNA synthesis was performed with random hexamer primers. Libraries were sequenced on an Illumina HiSeq2500 (4 libraries per lane). Reads were mapped to the mouse genome build NCBI37/mm9 using Bowtie2 and quantified using Seqmonk (Babraham Bioinformatics; http://www.bioinformatics.babraham.ac.uk/projects/seqmonk/). Differential expression analysis was performed using DESeq2 (Love et al., 2014), using all annotated genes, V_H_ genic transcripts and intergenic transcripts in a single analysis.

### VDJ-seq

VDJ-seq was performed as described (Bolland et al., 2016) for WT BM pro-B cells. VDJ-seq libraries for IL-7Rα^-/-^ pro-B cells were constructed similarly with two exceptions: firstly, due to reduced cell numbers, IL-7Rα^-/-^ VDJ-seq libraries were generated from 1.5-2 μg of DNA, compared to 10 μg of starting material used to generate the WT BM and FL libraries; secondly, due to the expectation for less variable VDJ events in IL-7Rα^-/-^ cells, measures were taken to distinguish and discount technical duplicates in VDJ-seq libraries. VDJ-seq libraries were de-duplicated based on the sequence of read 2 (containing both V_H_-DJ_H_ and D_H_-J_H_ junctions) and the sequence and position of the V_H_ gene (LinkON pipeline described in Bolland et al., 2016). This method relies on the variability of gene usage, junction diversity and the sonication step; therefore, reduced variability in the junctions and in V_H_ usage would lead to sequences being more likely to be identical, particularly in partially (D_H_-J_H_) recombined alleles. To overcome this issue, modified PE1 adaptors containing random barcodes were used to generate the IL-7Rα^-/-^ and FL libraries, allowing PCR duplicates, biological duplicates and Illumina sequencing errors to be distinguished (Chovanec et al., 2018). Libraries were sequenced on an Illumina HiSeq (8-12 libraries/lane; 250 BP paired-end). Reads were mapped to the mouse genome build NCBI37/mm9 using Bowtie2. To quantify individual V and D genes, probes were created over each gene segment and correctly orientated reads were quantified over each probe using Seqmonk. Libraries were also analysed using IMGT (International ImMunoGeneTics information system – http://www.imgt.org/ Lefranc et al., 2014). Analysis of VDJ-recombined sequences was carried out as described previously (Bolland et al., 2016); however, to analyse DJ-recombined sequences using IMGT, it was necessary to artificially add a V_H_ gene 5’ of the D_H_ sequence, as IMGT/HighVQUEST can only process VDJ sequences. The J558.78.182 V_H_ gene was appended, as it is functional and in-frame. These data were kept separate, and only D_H_ genes and the DJ_H_ junction were used for the analysis. Due to the low number of cells in IL-7Rα^-/-^ BM, VDJ-seq libraries were generated with approximately 6-fold less starting material, resulting in reduced numbers of sequences relative to the WT BM VDJ-seq libraries (STable 3). Nevertheless, this amount of starting material does not compromise detection of the wide dynamic range of frequency of VDJ and DJ recombined sequences (Chovanec et al., 2018).

### ATAC-seq

ATAC-seq (Assay for Transposase-Accessible Chromatin using sequencing) was performed as previously described (Buenrostro et al., 2013; 2015) on 70,000-100,000 cells. Sorted cells were washed, centrifuged and resuspended in 50 μl of lysis buffer (10 mM Tris-HCl pH 7.4, 10 mM NaCl, 3 mM MgCl2, 0.1% NP40) on ice for 15 min. Nuclei were centrifuged for 10 min at 4°C, resuspended in 50 μl 1xTD buffer containing 2.5 μl TDE1 transposase (Illumina Nextera DNA Sample Preparation Kit), and incubated for 30 min at 37°C. Samples were purified using MiniElute columns (Qiagen) according to the manufacturer’s instructions and eluted in 21 μl RSB buffer (10mM Tris HCl pH7.6, 10mM NaCl, 1.5mM MgCl2, 0.1% NP40). PCR amplification and index incorporation were performed in a 50 μl reaction containing 5 μl of forward and reverse index primers (Illumina Nextera Index Kit), 15 μl NPM, 5 μl PPC (Illumina Nextera DNA Sample Preparation Kit) and 20 μl DNA. Libraries were purified using QIAquick PCR clean-up columns (Qiagen) and sequenced on an Illumina HiSeq (6 libraries/lane). Reads were mapped to the mouse genome build NCBI37/mm9 using Bowtie2 and quantified using the MACS peak caller within Seqmonk. DESeq2 was used to identify genomic locations exhibiting significant differences in ATAC-seq reads in IL-7Rα/Rag2^-/-^ relative to Rag2^-/-^, and these sites were tested for TF motifs using the HOMER analysis tool (http://homer.salk.edu/homer/ngs/peaks.html - Heinz et al., 2010).

## Statistical test

A two-tailed ANOVA (type III) together with a pairwise t-test (adjusted by Benjamini and Hochberg method) were used to calculate significance and p-values unless otherwise stated.

## Data Availability

The VDJ-seq, ATAC-seq and RNA-seq raw sequencing files generated in this study, as well as processed files have been deposited with GEO, accession number GSE157603.

## Acknowledgments

We thank Geoff Butcher, Martin Turner and members of the Corcoran lab for helpful discussions and critical reading of the manuscript. We thank Dr Mark Veldhoen for provision of IL7Ra/Rag2 mice. Work in our Laboratory is supported by grants from the BBSRC (BBS/E/B/000C0404, BBS/E/B/000C0405, BBS/E/B/000C0427, BBS/E/B/000C0428, Core Capability Grant); A.B-E was supported by an MRC PhD studentship (1236141); B.A.S was supported by an MRC PhD studentship (1129229); M.J.T.S. was supported by a Babraham Institute PhD studentship.

## Author Contributions

Conceptualisation, A.B-E., B.A.S., and A.E.C.; Methodology, A.B-E., B.A.S, D.J.B.; Software, B.A.S., M.J.T.S., S.A; Validation, A.B-E., A.E.C.; Formal analysis, A.B-E., B.A.S., M.J.T.S., S.A.; Investigation, A.B-E., B.A.S.; Resources, A.B-E., K.T., A.E.C.; Data Curation, A.B-E., B.A.S.; Writing - Original Draft, A.B-E., A.E.C.; Writing - Review and Editing, A.B-E., B.A.S., M.J.T.S., D.J.B., S.A. A.E.C.; Visualization, A.B-E., B.A.S.; Supervision, A.E.C.; Project Administration, A.B-E. A.E.C.; Funding Acquisition, A.E.C.

