## Supplemental Figures for "IL-7R signalling activates widespread V_H_ and D_H_ gene usage to drive antibody diversity in bone marrow B cells"

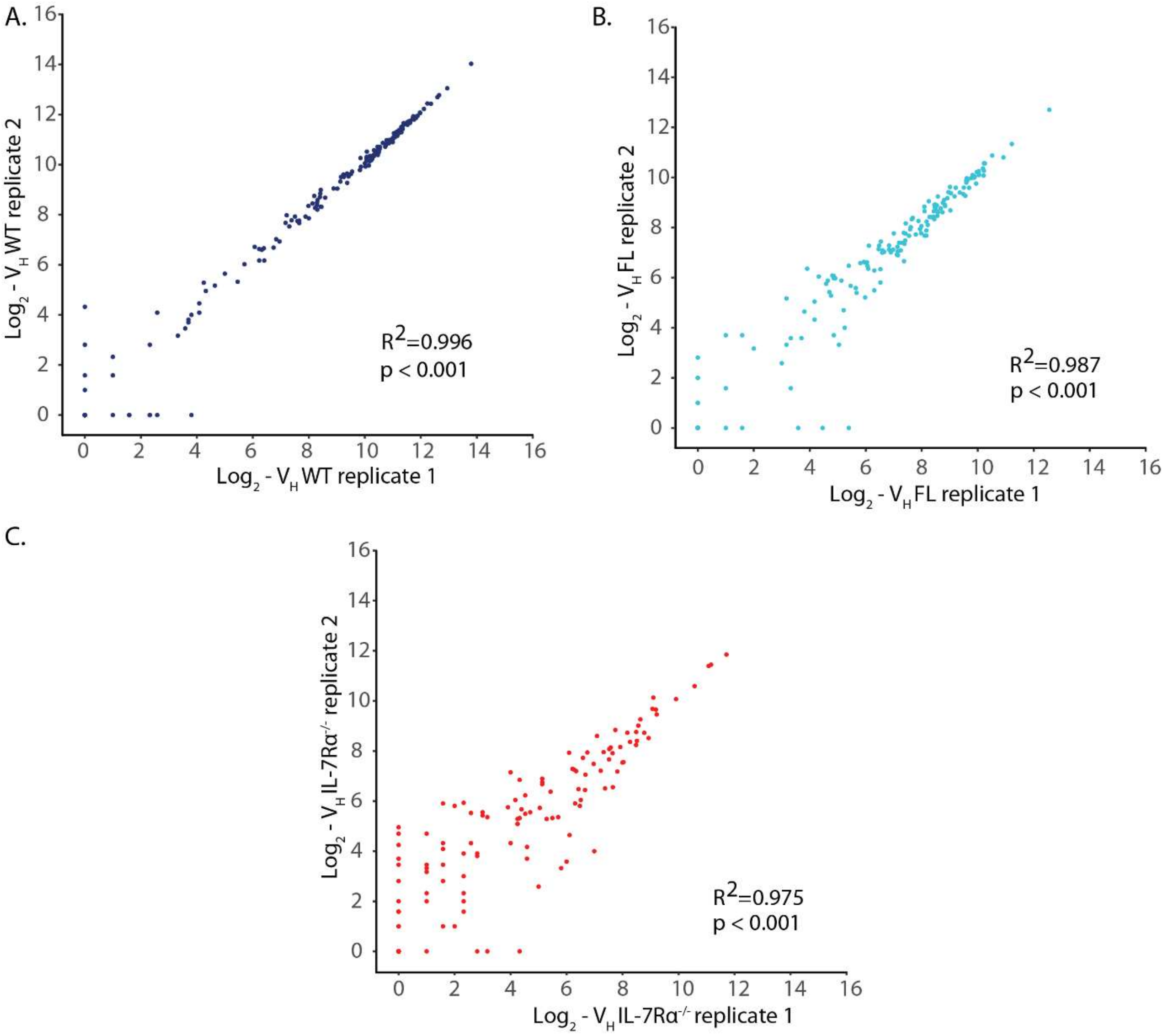


**S1: Replicate comparison between VDJ-seq replicates.** Scatterplot of VDJ-seq read counts (log2 transformed) of $V_H$ genes in two **A.** WT, **B.** FL and **C.** IL-7Rα$^{-/-}$ pro-B cell replicate datasets.

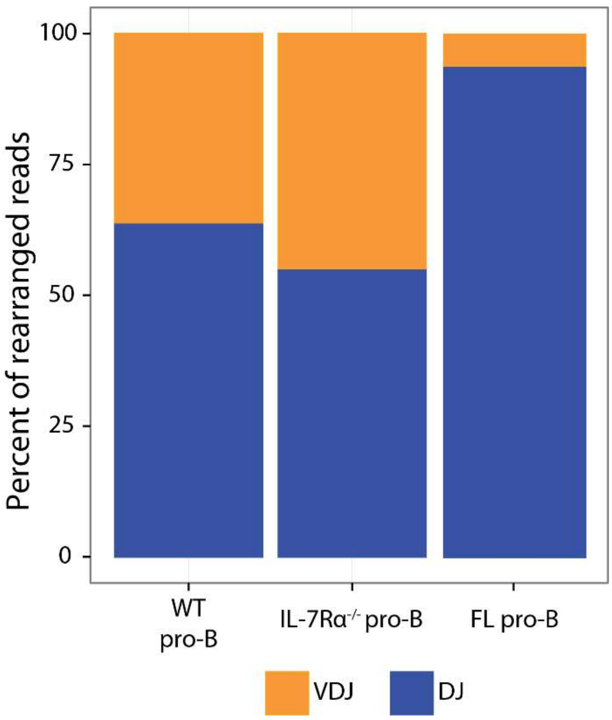


**S2: Proportion of $DJ_H$ and $VDJ_H$ sequences in each cell type.** The number of reads in the correct orientation was calculated over the $D_H$ and $V_H$ regions, and is shown as a percentage of total VDJ-seq reads in the correct orientation (mean of two replicates).

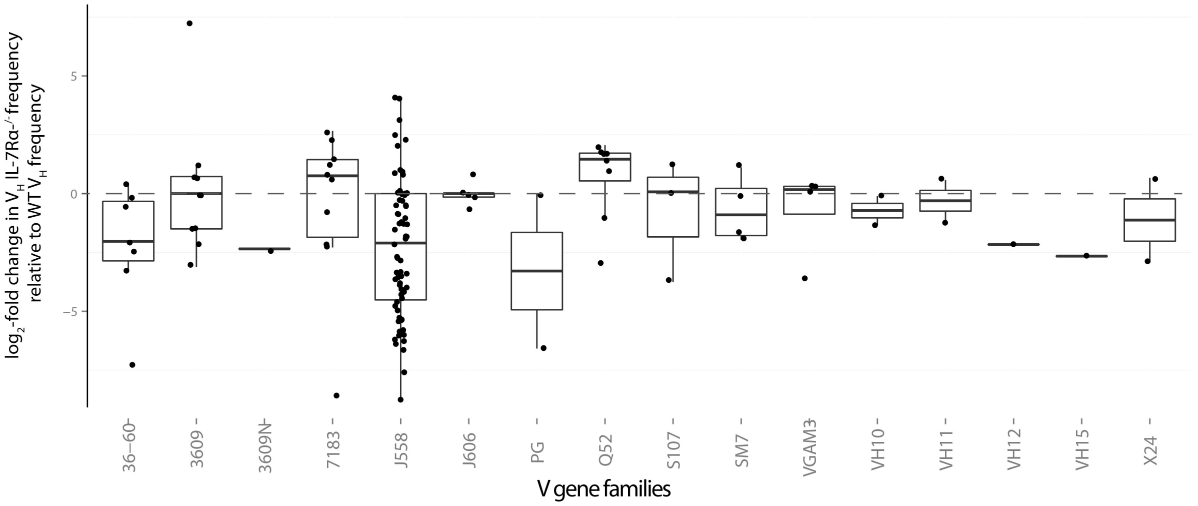


**S3: VDJ-seq values of each $V_H$ gene grouped in families.** The average of two WT and two IL-7Rα$^{-/-}$ VDJ-seq replicates was calculated for each $V_H$ gene. To display changes between WT and IL-7R-modified model frequencies, $V_H$ frequencies for each model were divided with the WT mean value and log2-transformed for comparison between models. Log2 values for each gene were grouped by gene family.

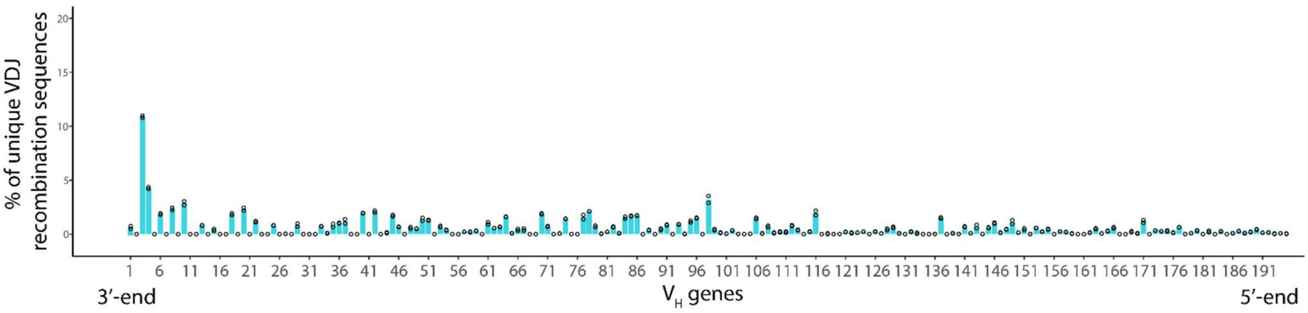


**S4: Fetal liver recombination.** Recombination frequencies of the 195 $V_H$ genes measured by VDJ-seq, for FL pro-B cells from 15.5 day old wild-type mice (50 embryos per replicate). Two replicates are shown as open circles. Reverse-strand reads were quantified for each $V_H$ gene and shown as a percentage of total number of reads quantified. For list of $V_H$ genes see supplementary table 1.

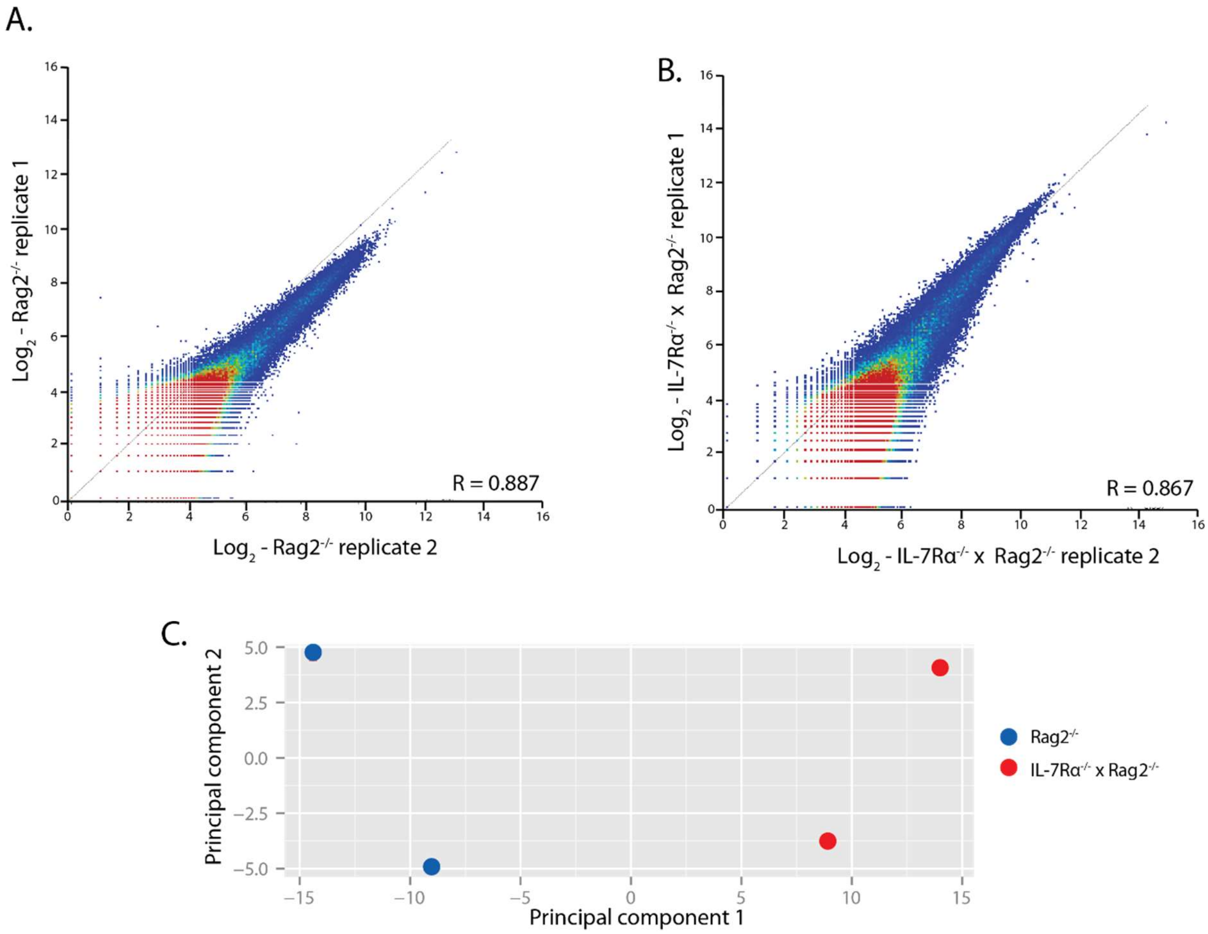


**S5: ATAC-seq libraries are highly correlated.** MACS peak caller was used to identify read peaks in both genotypes. ATAC-seq reads were quantified for peak probes for each replicate separately for **A.** Rag2$^{-/-}$ (Pearson's correlation R = 0.887) and **B.** IL-7Rα$^{-/-}$/Rag2$^{-/-}$ (R = 0.867) pro-B cell libraries. **C.** Principal component analysis show clustering of each genotype. Principal components are calculated based on the regularized-logarithm transformation of read counts per gene and using the R package pcaMethods. Analysis done using DESeq2 package.

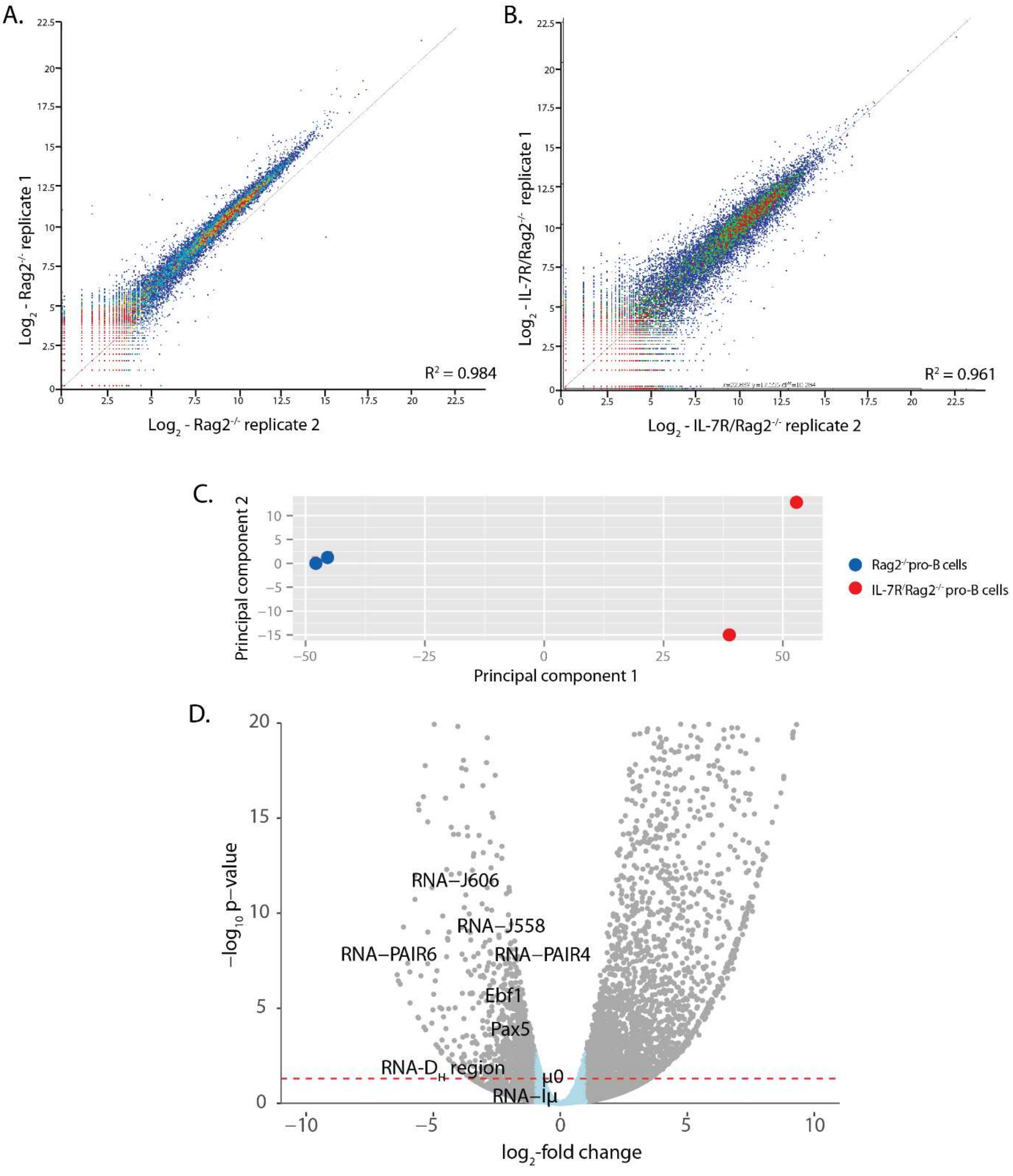


**S6: RNA-seq replicate libraries are highly correlated.** RNA-seq reads were quantified over all genes for each replicate for **A.** Rag2$^{-/-}$ and **B.** IL-7Rα$^{-/-}$/Rag2$^{-/-}$ pro-B cell libraries. **C.** Principal component analysis show clustering of each genotype. Principal components are calculated based on the regularized-logarithm transformation of read counts per gene and using the R package pcaMethods. Analysis done using DESeq2 package. **D.** Volcano plot showing genome-wide RNA-seq data. Grey dots are genes that are differentially expressed > 2-fold (in blue < 2-fold), dashed red line shows where p = 0.05. Transcripts of interest are tagged by name.

**S Table 1: Recombination activity of each $V_H$ gene determined by the binomial test.** VDJ-seq reads were quantified over each gene for each replicate library and raw read counts are shown for each replicate. A binomial test was used on the mean of two replicates to determine genes with significantly greater read counts than would be expected by chance, and were therefore considered to be actively recombining (R) and those which are not actively recombining (NR) in pro-B cells from WT and IL-7Rα$^{-/-}$ bone marrow cells, and in wild-type FL cells (fdr-adjusted p-value < 0.01).

**S Table 2: Quantification and analysis of transcription over $V_H$ gene segments in Rag2$^{-/-}$ and IL-7Rα$^{-/-}$/Rag2$^{-/-}$ pro-B cells.** RNA-seq raw reads were quantified over each $V_H$ gene probe (only $V_H$ genes which had detectable transcription are shown). Each of two replicates was quantified separately to determine differences in transcription between Rag2$^{-/-}$ and IL-7Rα$^{-/-/}$Rag2$^{-/-}$ pro-B cells. The adjusted mean for each probe, as well as the log2-fold change and adjusted p-value in the IL-7Rα$^{-/-}$/Rag2$^{-/-}$ relative to Rag2$^{-/-}$ samples was calculated by DESeq2. * - Significantly differentially expressed genes.

**S Table 3: VDJ-seq mapped read quantification.** VDJ-seq reads were mapped to the C57BL/6 mouse genome (genome build mm9). Paired-end read counts are shown for total mapped reverse orientated reads, and mapped reads over the whole $V_H$ and $D_H$ regions of the Igh locus. All samples were generated with 10 µg of pro-B cell DNA, except IL-7Rα$^{-/-}$ replicates which were made with 1-2 µg of DNA.

| Genotype | Total mapped reads | Reads mapped V region | Reads mapped D region |
|---|---|---|---|
| WT bone marrow 1 | 813k | 221k | 389k |
| WT bone marrow 2 | 898k | 240k | 449k |
| WT foetal liver 1 | 757k | 55k | 643k |
| WT foetal liver 2 | 920k | 62k | 785k |
| IL-7Rα$^{-/-}$ bone marrow 1 | 57k | 21k | 27k |
| IL-7Rα$^{-/-}$ bone marrow 2 | 71k | 26k | 30k |
